# Structural differences between pri-miRNA paralogs promotes alternative Drosha cleavage and expands target repertoires

**DOI:** 10.1101/366492

**Authors:** Xavier Bofill-De Ros, Wojciech K. Kasprzak, Yuba Bhandari, Lixin Fan, Quinn Cavanaugh, Minjie Jiang, Lisheng Dai, Acong Yang, Tie-Juan Shao, Bruce A. Shapiro, Yun-Xing Wang, Shuo Gu

## Abstract

MicroRNA (miRNA) processing begins with Drosha cleavage, the fidelity of which is critical for downstream processing and mature miRNA target specificity. To understand how pri-miRNA sequence and structure influence Drosha cleavage, we studied the maturation of three pri-miR-9 paralogs, which encode the same mature miRNA but differ in the surrounding scaffold. We show that pri-miR-9-1 has a unique Drosha cleavage profile due to its distorted and flexible stem structure. Cleavage of pri-miR-9-1, but not pri-miR-9-2 or pri-miR-9-3, generates an alternative-miR-9 with a shifted seed sequence that expands the scope of its target RNAs. Analyses of low grade glioma patient samples indicate that the alternative-miR-9 plays a distinct role in preventing tumor progression. To generalize our model, we provide evidence that distortion of pri-miRNA stems correlates with Drosha cleavage at non-canonical sites. Our studies reveal that pri-miRNA paralogs can have distinct functions via differential Drosha processing.

## Introduction

MicroRNAs (miRNAs) are a class of small non-coding RNAs (~22nt) that are highly conserved among species (Bartel, 2018; Pasquinelli, 2012). Collectively, miRNAs negatively regulate the expression of >60% of human genes by post-transcriptional mechanisms (Friedman et al., 2009). It is well-established that miRNAs play critical roles in the development of many diseases, including cancer (Lin and Gregory, 2015). Under the canonical biogenesis pathway, miRNAs are transcribed as long primary transcripts (pri-miRNAs), the maturation of which requires stepwise cleavage by two RNase III enzymes, Drosha and Dicer (Ha and Kim, 2014). Drosha and its cofactor DGCR8 form the Microprocessor complex in the nucleus, cutting the pri-miRNA to release the precursor miRNA (pre-miRNA) in the form of a short hairpin (Denli et al., 2004; Gregory et al., 2004; Han et al., 2004; Lee et al., 2003). The pre-miRNA is exported to the cytoplasm, where it is processed further by Dicer (Grishok et al., 2001; Hutvágner et al., 2001; Knight and Bass, 2001). The resulting duplex is loaded onto one of four Argonaute proteins (AGO1-4), forming the RNA-induced silencing complex (RISC). During this process, one of the two strands (miRNA or guide strand) is retained while the other (passenger strand) is discarded (Khvorova et al., 2003; Schwarz et al., 2003). In mammals, most miRNAs are partially complementary to their targets, inducing translational repression and/or mRNA decay (Béthune et al., 2012; Gu and Kay, 2010; Iwakawa and Tomari, 2015; Jonas and Izaurralde, 2015). The nucleotides at positions 2 to 7, counting from the miRNA 5’ end, are termed the seed region. Base-pairing of this region with its mRNA target is essential and sufficient for many miRNAs to function (Bartel, 2009).

The miRNA biogenesis machinery differentiates a wide variety of pri-miRNA hairpins from numerous other RNAs folded into similar structures. The pri-miRNA hairpin is structurally defined by the presence of a terminal loop, a stem of roughly three helical turns and basal unpaired flanking sequences. The stem is divided into an upper stem where the mature miRNA sequence is embedded and a lower stem next to the flanking region (T. A. Nguyen et al., 2015). Drosha senses the basal junction between the lower stem and the single-stranded basal region, establishing a cleavage site 11bp away (Han et al., 2006). In parallel, DGCR8 contacts the apical junction between the upper stem and terminal loop to facilitate proper recognition of pri-miRNA (T. A. Nguyen et al., 2015; Zeng et al., 2005). Recent findings indicate that certain sequence motifs in the pri-miRNA scaffold also contribute to efficient processing of pri-miRNA in mammalian cells (Auyeung et al., 2013; Fang and Bartel, 2015).

Although both Drosha and Dicer cleavages define the sequence of mature miRNAs, the former is more critical because it defines the latter: Dicer cleaves at a fixed distance from the terminal ends of pre-miRNA (MacRae et al., 2007; Park et al., 2011); Its cut site is largely determined by the Drosha cleavage site. Drosha cleavage at an alternative position generates a miRNA isoform (isomiRs) with a distinct 5’ end and an altered seed sequence. As these isomiRs will have altered target specificity, alternative cleavage by Drosha can profoundly impact miRNA function. Several studies have shown that changes in the relative distances between the expected cleavage site, the basal junction, and the apical junction of a pri-miRNA can all affect Drosha cleavage fidelity (Burke et al., 2014; Ma et al., 2013; Roden et al., 2017). However, despite the importance of Drosha cleavage in dictating miRNA specificity, the mechanisms by which Drosha selects its precise site of cleavage are largely unknown.

A class of pri-miRNAs that may be particularly susceptible to alternate cleavage by Drosha are those encoded by multigene families. Approximately 30-40% of miRNAs are encoded by multiple loci that derive from gene duplications. Many of the miRNAs encoded by more than one loci are predicted to have identical seed sequences and overall homology (Berezikov, 2011). One example are the miRNAs encoded by the human pri-miR-9 family, which consists of three members encoded on chromosome 1 (pri-miR-9-1), chromosome 5 (pri-miR-9-2) and chromosome 15 (pri-miR-9-3). Despite the complete conservation of the mature miR-9 sequence within the upper stems of all three pri-miRNAs, variations occur at other positions that could potentially affect biogenesis. MiR-9 is highly expressed in brain, where it is involved in neural stem cell self-renewal and differentiation (Zhao et al., 2009), and synaptic plasticity (Sim et al., 2016). Mir-9 levels are also dysregulated in many human cancers (Ma et al., 2010; Nowek et al., 2018). Consistent with a role for Drosha cleavage fidelity in driving function, an isomiR of miR-9 with an altered 5’ end was detected in human embryonic stem cells and neural cells and shown to affect target site selection of several reporter constructs (Tan et al., 2014.). However, both the mechanism(s) by which this isomer arises and its biological roles are unknown.

Here, we systematically investigate the biogenesis of miR-9 and its isomers. We report that although canonical miR-9 is derived from all three pri-miRNAs, the miR-9 5’ isomiR is primarily generated from pri-miR-9-1. We show that the distorted and flexible structure of the pri-miR-9-1 lower stem promotes Drosha cleavage at an alternative site, resulting in isomiR generation. Analysis of brain tumors transcriptomes suggests that the new isomiR downregulates specific targets to potentially regulate tumor progression. We provide evidence that non-canonical Drosha cleavage of other pri-miRNAs may similarly be driven by distortion of the pri-miRNA stems. Together, our findings reveal the pri-miRNA structural features that alter the fidelity of Drosha cleavage and provide insights into the mechanisms that result in neofunctionalization of miRNA paralogs.

## Results

### Pri-miR-9-1 generates a 5’ isomiR due to its unique Drosha cleavage profile

To investigate whether all three pri-miR-9 paralogs can produce mature miR-9, we first used a previously established reporter system to measure the efficacy of Drosha cleavage for each paralog (Dai et al., 2016). Briefly, three luciferase reporter constructs harboring each paralog sequence in their 3’UTR were co-transfected into Drosha knockout cells with either a Drosha-expressing vector or a control plasmid. In this assay, Drosha cleavage of the pri-miRNA sequence triggers degradation of the luciferase transcripts and reduces luciferase activity (Dai et al., 2016). Drosha cleavage efficiency can thus be measured by comparing reporter activity in the presence or absence of Drosha (Figure 1–figure supplement 1A). The three pri-miR-9 paralogs on the reporter undergo Drosha processing with similar efficiencies (Figure 1A), indicating that they are all recognized by Drosha.

To further examine maturation, we cloned the genomic sequence of each pri-miR-9 paralog and its surrounding sequence into a CMV (Pol II) driven transcription vector. By separately transfecting these plasmids into HEK293T cells, we were able to express each pri-miR-9 paralog independently. Northern blot analysis confirmed the production of mature miR-9 (hsa-miR-9-5p/MIMAT0000441) from all three pri-miR-9 transcripts (Figure 1B). Interestingly, multiple miRNA products varying in length were identified in all cases, suggesting alternative Drosha/Dicer cleavages and/or post-cleavage modifications. Endogenous miR-9 was barely detectable (Figure 1B), indicating that its abundance is marginal in these cells compared to the overexpressed miR-9 and therefore could be neglected in subsequent analyses.

To study the processing in detail, we deep-sequenced all miRNAs from the transfected cells and mapped the reads back to the corresponding pri-miR-9 paralog. Interestingly, while all pri-miR-9 transcripts produce the expected canonical miR-9 sequence (miR-9-can), an alternative miR-9 sequence (miR-9-alt) that begins one nucleotide downstream of the miR-9-can is generated exclusively from pri-miR-9-1 (14% of total reads) (Figure 1C). Consistent with Northern blot results (Figure 1B), both miR-9-can and miR-9-alt are heterogeneous in length due to their 3’ tailing and trimming as described previously (Berezikov et al., 2011; Gu et al., 2012). Similar results were obtained when pri-miR-9 paralogs were expressed in HeLa cells: miR-9-alt (6% of the total reads) is generated from pri-miR-9-1, but not from pri-miR-9-2 or pri-miR-9-3, indicating that our observations are not limited to a single cell type (Figure 1–figure supplement 1B,C).

Given that post-maturation sequence modifications at the 5’ of miRNA are extremely rare (Burroughs et al., 2010), the production of miR-9-alt is likely a result of Drosha cleavage at an alternative site. In fact, previous studies have established that the 5’ end of mature miRNA is a faithful indicator of the Drosha cleavage site (Gu et al., 2012; Ma et al., 2013). To test this directly, we performed an *in vitro* Drosha cleavage assay on pri-miR-9-1 and pri-miR-9-2 transcripts (Figure 1D). Consistent with the *in vivo* results, Drosha processing of pri-miR-9-1, but not pri-miR-9-2, generates an additional pre-miR-9 product shorter than the expected size, indicating the usage of an alternative cleavage site downstream of the canonical one. Moreover, data mining of a recent study in which Drosha cutting sites were mapped by sequencing the endogenous cleavage products (pre-miRNAs) crosslinked to Drosha (Kim et al., 2017) revealed that some pre-miR-9-1 RNAs, but not pre-mir-9-2 RNAs contained the alternative Drosha 5’ cleavage site (Figure 1–figure supplement 1D). Together, these results indicate that the 5’ isomiR miR-9-alt is generated from pri-miR-9-1 due to a unique Drosha cleavage profile that differs from the other two pri-miR-9 paralog transcripts.

**Figure 1.**
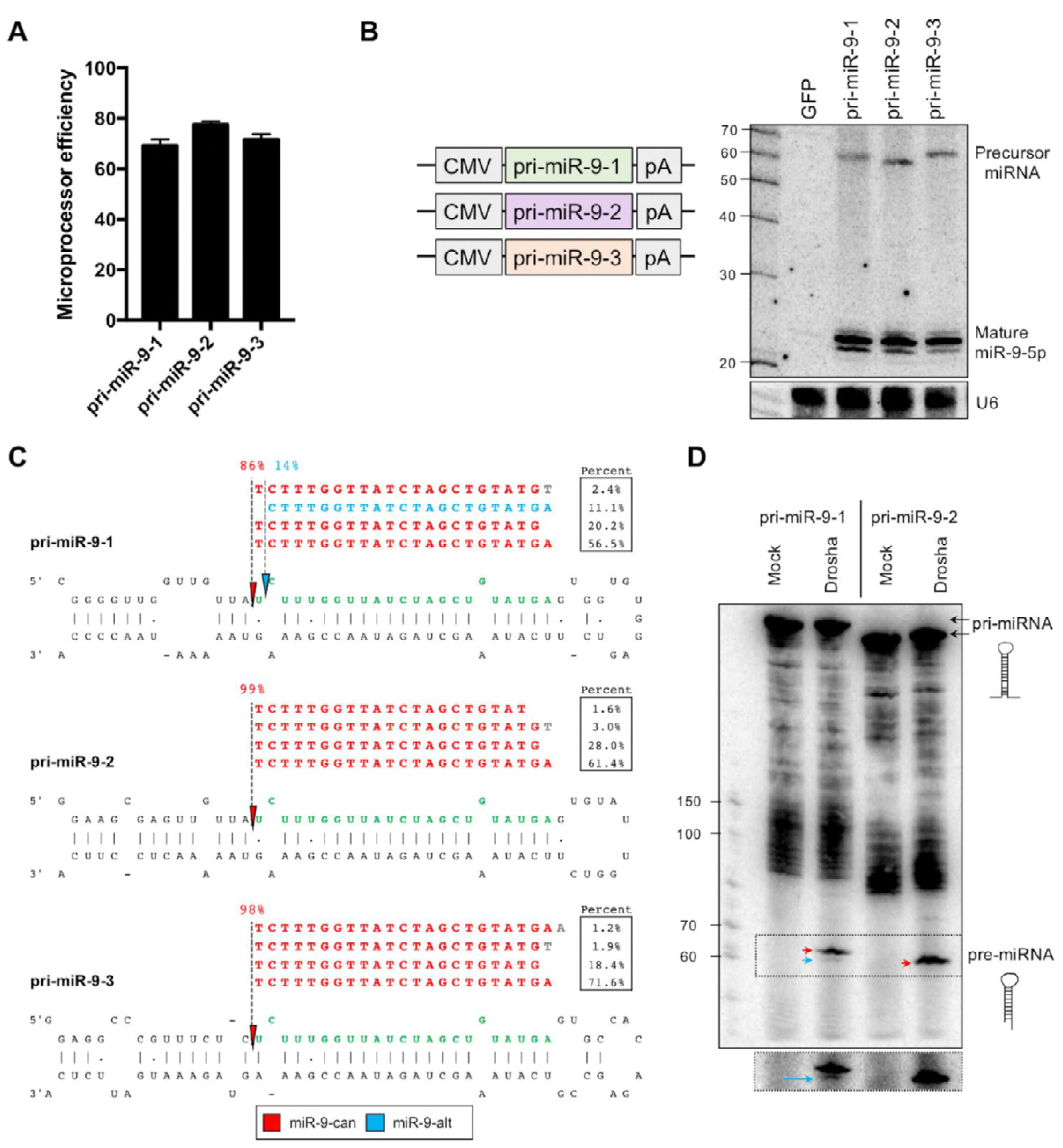
Pri-miR-9-1 has a unique Drosha cleavage profile. (***A***) *Renilla* luciferase reporters containing pri-miR-9 family members in the 3’UTR were transfected into HEK293T Drosha KO cells with or without the co-expression of Drosha. Dual-luciferase assays were performed 48h post-transfection. *Renilla* luciferase activities were normalized to Firefly luciferase activities and Microprocessor efficiency was calculated as described in the Methods section. Error bars represent the standard error of the mean (SEM) of three biological replicates. (***B***) CMV-driven pri-miR-9 cassettes were expressed individually in HEK293T cells. pre-miR-9 and mature miR-9 were detected by Northern blotting with probe-miR-9 (Supplemental Table S2). U6 snRNA was detected as an internal control. Note, pre-miR-9-2 (59nt) is shorter than pre-miR-9-1 (61nt) and pre-miR-9-3 (61nt) due to differences in the loop sequence. (***C***) Small RNAs from HEK293T cells transfected with pri-miR-9-expressing plasmids were subjected to deep sequencing. After being mapped to the corresponding pri-miR-9 paralog, only the four most abundant sequences were labeled in the figure along with their percent abundance relative to all pri-miR-9 reads. The percentage of sequences starting at a position relative to the total number of miR-9 reads was used to infer the Drosha cleavage percentage, which was labeled with a small solid arrow and a dotted line. miR-9-can and its tailed or trimmed isomiRs are in red; miR-9-alt is in blue. (***D***) *In vitro* transcribed pri-miR-9-1 or pri-miR-9-2 transcripts were incubated with mock or purified Microprocessor complex. Precursor miRNAs (pre-miR-9-1 and pre-miR-9-2) were detected by Northern blot using probe-miR-9 and indicated by a red arrow. A shorter product (indicated by a blue arrow) was only observed with pre-miR-9-1 but not with pre-miR-9-2. A longer exposure for a better visualization of the pre-miRNAs is also presented.

### miR-9-alt regulates a novel set of targets to repress low grade glioma (LGG)

The current model proposes that the seed region plays a critical role in defining miRNA target specificity (Bartel, 2009). Although miR-9-alt and miR-9-can originate from the same parental transcript (pri-miR-9-1), they have different seed sequences and therefore should each repress a unique set of targets. To evaluate this experimentally, we sought to decouple their biogenesis by expressing miR-9-can and miR-9-alt independently. We achieved this goal by constructing two well-designed U6 (pol III)-driven shRNAs (sh-miR-9-can and sh-miR-9-alt). Deep sequencing confirmed that each shRNA generates the corresponding mature miRNA isomiR with a negligible amount (<0.1%) of the other (Figure 2–figure supplement 1A). Luciferase reporters containing artificial target sites complementary to the seed sequence of either miR-9-can or miR-9-alt were co-transfected with these two shRNAs. As expected, sh-miR-9-can and sh-miR-9-alt only repress efficiently their corresponding target, establishing that we can measure the function of miR-9-can and miR-9-alt separately (Figure 2A). Seed-based prediction identified 539 unique targets of miR-9-alt, thus increasing the number of potential targets of miR-9 paralogs by over 57% (Figure 2B). Using luciferase reporters, we validated three known targets of miR-9-can (FOXG1, GABRB2 and HES1)(Bonev et al., 2012; Pietrzykowski et al., 2008; Shibata et al., 2008) as well as three novel targets of miR-9-alt (CSGALNACT1, FOXN3 and PURB)(Figure 2–figure supplement 1B). Together, these results demonstrate that miR-9-alt has the potential to regulate a distinct set of target genes in cells.

To identify potential consequences of mir-9-alt expression, we searched datasets for cells and tissues in which this miRNA was well-expressed. Consistent with previous reports describing the relevance of miR-9 in neuronal development and homeostasis (Coolen et al., 2013), we found that low-grade glioma (LGG) has the highest average expression of miR-9 among tumors documented in The Cancer Genome Atlas (TCGA) (Figure 2–figure supplement 1C). Further analysis revealed that the abundance of miR-9-alt, albeit only ~10% of miR-9-can, is still higher than the levels of miRNAs with well-established functions such as let-7, miR-30 or miR-21 (Figure 2C). To investigate whether miR-9-alt regulates predicted miR-9-alt-specific targets in LGG, we took advantage of the RNA-Seq and miRNA-Seq data in TCGA. Specifically, we compared mRNA expression profiles between patients with high levels of miR-9-alt (top 5%, 25 samples) and relatively low levels of miR-9-alt (bottom 5%, 25 samples) (Figure 2–figure supplement 1D). As expected, those mRNAs with one predicted target site of miR-9-alt are repressed in comparison to the whole transcriptome. Increased repression was observed with mRNAs containing multiple target sites, suggesting the repression is specific to miR-9-alt (Figure 2D). Applying 2-fold repression as a threshold, we identified 40 genes that are subject to miR-9-alt regulation (Supplemental Table S1).

Next, we sought to interrogate the biological impact of miR-9-alt in LGG. Gene ontology analysis (GO term) revealed an enrichment of extracellular matrix remodeling factors among the identified miR-9-alt targets. Many of these factors, including BACE2, COL1A2 and FGL2, have documented roles in promoting tumorigenesis (Balbous et al., 2014; Liu et al., 2010; Oliveras-Ferraros et al., 2014; Shin et al., 2017; Yan et al., 2015). Consistent with this idea, we found that lower levels of BACE2, COL1A2 and FGL2 correlate with increased survival time for the LGG patients (Figure 2E). Results of luciferase reporter assays confirmed that these targets are specifically inhibited by miR-9-alt (Figure 2F), suggesting miR-9-alt could function as a tumor suppressor. The host genes of pri-miR-9-1 and pri-miR-9-2 are highly expressed (FPKM > 10) in LGG (Figure 2–figure supplement 1E) and both contribute to the production of miR-9-can. miR-9-alt, on the other hand, originates exclusively from pri-miR-9-1 (Figure 1C). In support of a tumor suppressor role of miR-9-alt, the level of pri-miR-9-1, but not pri-miR-9-2, is positively correlated with LGG patient survival with statistical significance (p<0.01) (Figure 2G; Figure 2–figure supplement 1F).

Taken together, these results indicate that miR-9-alt has a biological impact on the progression of LGG, most likely through inhibiting BACE2, COL1A2 and FGL2. Moreover, we also identified the glutamate receptor NMDA2A as a target of miR-9-alt (Figure 2–figure supplement 1G). Thus, miR-9-alt likely has functions beyond its role in tumor progression and may affect brain physiology.

**Figure 2.**
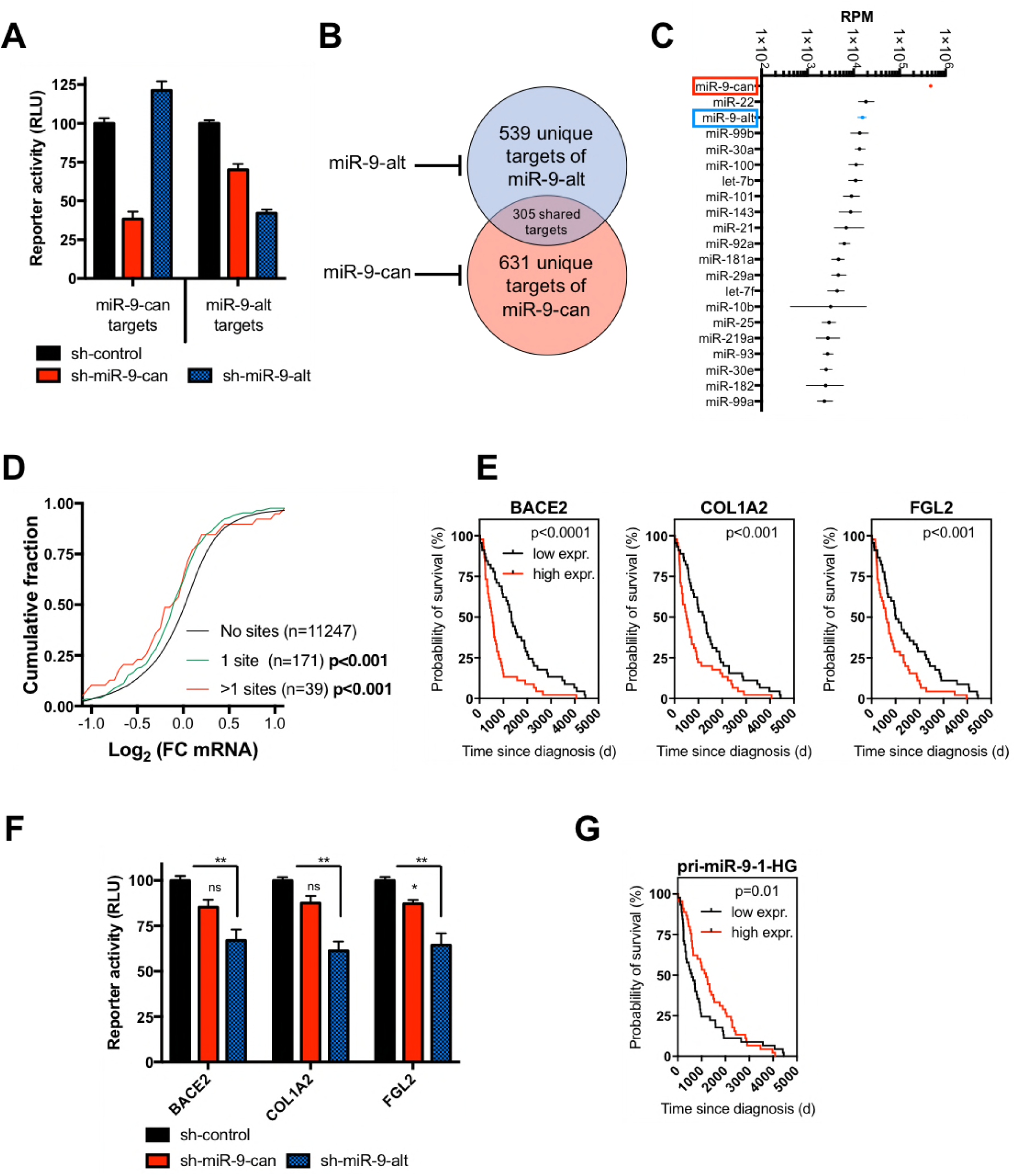
miR-9-alt regulates a novel set of targets to repress low grade glioma (LGG). (***A***) Repression of miR-9 isomiRs was measured by dual-luciferase reporter assay. The psi-CHECK2 vector with two tandem target sites in the 3’ UTR and DNA plasmids expressing shRNA were co-transfected into HEK293 cells. These target sites were designed with a 7mer-A1 seed region plus additional base-pairs on the 3’ end of the miRNA. Dual-luciferase assays were performed 48h post-transfection. *Renilla* luciferase activities were normalized with firefly luciferase, and the percentage of relative enzyme activity compared to the negative control (treated with sh-control) was plotted. Error bars represent the SEM from three biological replicates. (***B***) Venn diagram of predicted targets of miR-9-can and miR-9-alt by TargetScan (Lewis et al., 2005). (***C***) Average expression level (reads per million, RPM) of miR-9-can and miR-9-alt among top expressed miRNAs in LGG tumors. Error bars represent the standard deviation (SD) over 525 LGG patient samples documented in TCGA. (***D***) Cumulative fraction plot of fold-change in expression of mRNAs between the top and low levels of the miR-9-alt in patients from LGG. Shift towards the left indicates repression. Also see Figure 2–figure supplement 1 for details. (***E***) Kaplan-Meier survival curve of LGG patients with high (red line) and low (black line) levels of the target gene. (***F***) Predicted miR-9-alt target sites from each gene were cloned individually in the 3’UTR of the luciferase reporter. Dual-luciferase assays were performed at 48h post-transfection, and the results were plotted as described previously. Error bars represent the SEM from three biological replicates. *, p<0.05 **, p < 0.01 n.s., non-significant. **(*G*)** Kaplan-Meier survival curve of LGG patients with high (red line) and low (black line) levels of the pri-miR-9-1 host gene. p-values were calculated using Wilcoxon test.

### The lower stem plays a major role in determining cleavage fidelity

Next, we sought to understand why miR-9-alt is generated exclusively from pri-miR-9-1. In these experiments, we investigated why Drosha processes pri-miR-9-1 and pri-miR-9-2 differently. The pri-miR-9-1 and pri-miR-9-2 scaffolds differ in three regions: the terminal loop (TL), the lower stem (LS), and the 5’ and 3’ flanking sequences (F) (Figure 3A). To determine which element drives the changes on Drosha cleavage fidelity, we swapped these regions between pri-miR-9-1 and pri-miR-9-2, generating six chimeric pri-miRNAs. These chimeric constructs were expressed individually in HEK293T cells. Drosha cleavage efficiency was measured by the reporter assay and Northern blotting. In parallel, cleavage fidelity was determined by deep sequencing. While all chimeras were processed with similar efficiency (Figure 3–figure supplement 1A,B), the fidelity of cleavage varied. The alternative cleavage is nearly abolished when the lower stem of pri-miR-9-1 is replaced with that of pri-miR-9-2 (F1-LS2-TL1) whereas exchanging the loop (F1-LS1-TL2) or the flanking sequences (F2-LS1-TL1) only partially reduces usage of the alternative cleavage site (Figure 3B; Figure 3–figure supplement 1C). Consistent with the finding that features of the lower stem are important for fidelity, a pri-miR-9-2 scaffold in which only the lower stem is replaced by that of pri-mir-9-1 stem (F2-LS1-TL2) shows an increase in the rate of alternative Drosha cleavage. In contrast, a pri-mir-9-2 scaffold containing either the pri-mir-9-1 loop (F2-LS2-TL1) or pri-mir-9-1 flanking sequences (F1-LS2-TL2) did not give increased cleavage at the alternative site (Figure 3C; Figure 3–figure supplement 1D). Thus, the lower stem is the major determinant for inducing alternative cleavage of pri-miR-9-1.

Similar results were obtained when these chimeras were expressed in HeLa cells (Figure 3–figure supplement 1E,F). Taken together, our results demonstrate that the lower stem of pri-miR-9-1 is both necessary and sufficient for the increased level of Drosha alternative cleavage. Interestingly, the lower stem sequence of pri-miR-9-1 is well conserved in vertebrates. In contrast, the lower stem sequences of pri-miR-9-2 and pri-miR-9-3 as well as the terminal loop sequences of all pri-miR-9 paralogs are variable (Figure 3–figure supplement 1G). This suggests that Drosha alternative cleavage of pri-miR-9-1 and therefore the production of miR-9-alt may be evolutionarily conserved.

**Figure 3.**
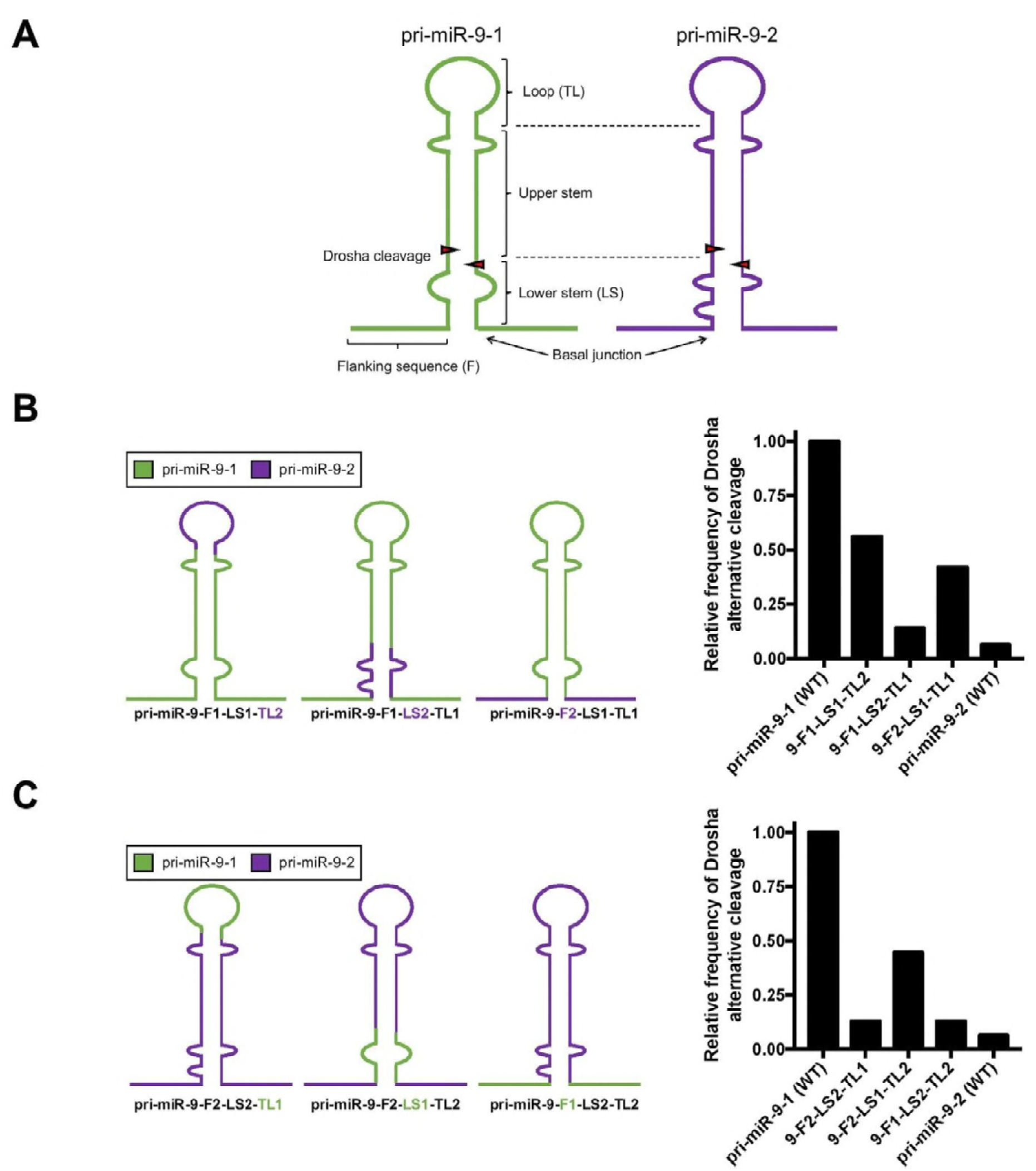
The lower stem plays a major role in determining cleavage fidelity. (***A***) Schematic representation of pri-miRNA structural elements. (***B***) Left panel, schematic representation of chimeric constructs where structural elements of pri-miR-9-1 have been replaced for their equivalents in pri-miR-9-2. Note that flanking sequences (F), lower stem (LS) and terminal loop (TL) from pri-miR-9-1 are depicted in green while these from pri-miR-9-2 are in purple. Right panel, all chimeras were expressed in HEK293T cells. Relative Drosha alternative cleavage frequencies on each chimeric construct were measured by deep sequencing and plotted here. (***C***) Left panel, schematic representation of chimeric constructs where structural elements of pri-miR-9-2 have been substituted for their equivalents in pri-miR-9-1. Right panel, relative Drosha alternative cleavage frequencies on each chimeric construct in HEK293T cells. Note, in both (***B,C***), Drosha alternative cleavage frequency on wild-type pri-miR-9-1 (14%) was set as 1 in the figure.

### The distorted lower stem structure drives alternative Drosha cleavage

The current model proposes that Drosha cleavage fidelity is mainly determined by the length of the lower stem. Specifically, Drosha prefers to cleave at a 5’ site 13nt away and a 3’ site 11nt away from the basal junction (Auyeung et al., 2013; Han et al., 2006; Ma et al., 2013; T. A. Nguyen et al., 2015). Consistent with this model, we found that the pri-miR-9-1 lower stem is a major determinant for Drosha cleavage fidelity. However, none of the pri-miR-9 paralogs’ lower stem lengths are optimal (Figure 4–figure supplement 1A), suggesting that additional factors underlie the Drosha alternative cleavage of pri-miR-9-1. We speculated that structural features might contribute to the choice of Drosha cleavage site. Indeed, secondary structure predictions revealed that pri-miR-9-1 contains an internal loop near the cut site (Figure 4–figure supplement 1A). However, pri-miR-9-2 and pri-miR-9-3 also contain bulges and mismatches in their lower stems (Figure 4–figure supplement 1A), making an internal loop per se an unlikely cause of Drosha alternative cleavage.

To further investigate the structural differences between the three paralogs, we analyzed their tertiary structures with RNAComposer (Popenda et al., 2012). These initial predicted structures were then refined using molecular dynamics. In brief, the position and motion of each atom were calculated every 2 femtoseconds over the course of 250 nanoseconds. Interestingly, while pri-miR-9-2 and pri-miR-9-3 are predicted to maintain a relatively straight helix during the course of simulation, tertiary structure of pri-miR-9-1 is predicted to be distorted and flexible. The folding structure of pri-miR-9-1 with lowest energy is bent at lower stem (Figure 4–figure supplement 1B). To validate these predicted topologies, we used small angle X-ray scattering (SAXS). SAXS is a solution-based method that does not require crystallization and provides information on overall molecular size, shape, intermolecular distances and dynamics (Fang et al., 2013). We used ensemble calculations to characterize small and large amplitude motions of the three-dimensional structure of the paralog stems. As predicted, the lower stem of pri-miR-9-1 is kinked whereas that of pri-miR-9-2 is not (Figure 4A,B; Figure 4–figure supplement 1C).

Since the sequence of the lower stem is conserved in vertebrates (Figure 3–figure supplement 1G), we tested whether the sequence of the bulge was important by creating two additional pri-miR-9-1 constructs (pri-miR-9-1-a and pri-miR-9-1-b) with changes in the lower stem bulge nucleotide composition (Figure 4C). Interestingly, while both pri-miR-9-1-a and pri-miR-9-1-b have the same secondary structure as that of pri-miR-9-1 (a 4nt by 3nt asymmetric bulge), their 3D predicted structures and dynamics are different. In fact, multiple structures with variable degree of distortion at the lower stem were generated for both pri-miR-9-1-a and pri-miR-9-1-b, indicating their tertiary structures are highly flexible (Figure 4C). Correspondingly, the frequency of Drosha cleavage at the alternative site is much higher (54% and 83% respectively) in HEK293T cells and similar results were obtained in HeLa cells (Figure 4C; Figure 4–figure supplement 1D,E). These results suggest that the distorted and flexible lower stem structure, which is apparently a result of the asymmetrical bulge, induces alternative Drosha cleavage. To test this, we sought to correct the distortion and flexibility of the lower stem of pri-miR-9-1 without changing its length (Figure 4–figure supplement 1F). We achieved this goal by either replacing the bulge with a perfect stem (pri-miR-9-1-perfect-stem) or by adding a single nucleotide to the bulge (pri-miR-9-1-bulged-stem) (Figure 4D). Molecular dynamics modeling confirmed that these modified structures form a relatively rigid and straight helix at the lower stem (Figure 4–figure supplement 1G). Deep sequencing analysis revealed that both of the modified pri-miR-9-1s are nearly free of Drosha-mediated alternative cleavage, instead resulting in a cleavage profile similar to that of pri-miR-9-2 in both HEK293T and HeLa cells (Figure 4D; Figure 4–figure supplement 1H,I). These results demonstrate that a 3D structural feature of pri-miRNA, specifically a distorted and flexible stem, impacts Drosha cleavage fidelity.

**Figure 4.**
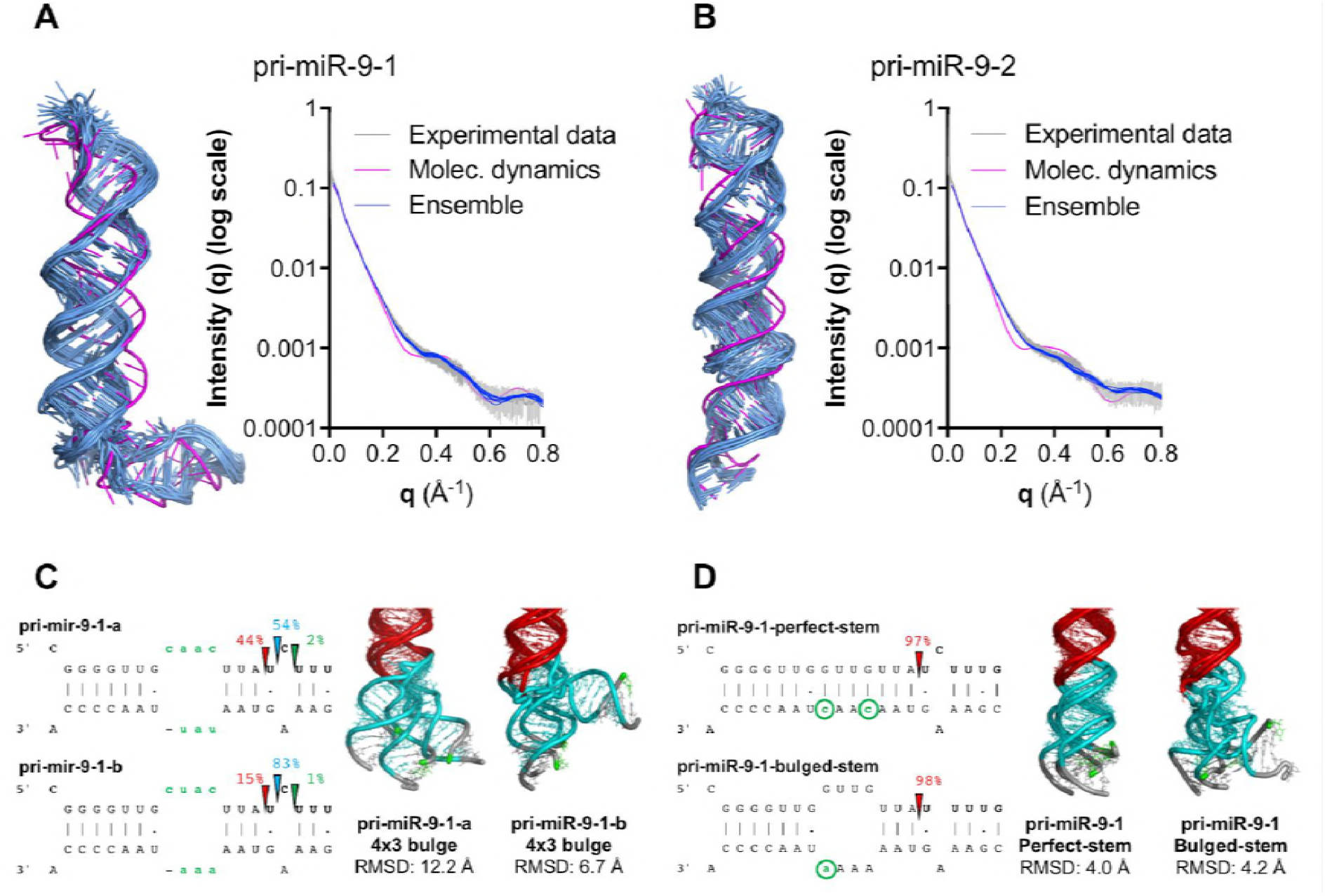
The tertiary structure of the lower stem drives alternative Drosha cleavage. (***A,B***) Left, three dimensional structures of pri-miR-9-1 and pri-miR-9-2 obtained after molecular dynamics refinement (magenta) and top ten ensemble structures derived from the SAXS data (blue). Right, experimental SAXS data (gray line) plotted as scattering intensity (arbitrary units) versus momentum transfer q (Å^−1^). The plot also displays the calculated SAXS curves of the molecular dynamics structure (magenta line) and the Intensity (q) values obtained in the top ten ensemble structures (blue lines). As one can see SAXS curves are very sensitive to the shape of the molecule and the ensemble calculations give rise to the best fit to the experimental SAXS data. (***C,D***), Left, secondary structure of the lower stem of pri-miR-9-1 mutants. Green labeling illustrates the nucleotides changed in each case. The arrows and the corresponding numbers indicate the inferred Drosha cleavage sites and their relative percentage. Right, local tertiary structure motions of pri-miR-9-1 mutants: lower stem (cyan); upper stem (red). The distortion is evaluated by RMSD (root-mean-square deviation) during molecular dynamics against the backbone of a perfect RNA A-form helix.

### Pri-miRNA tertiary-structure-based alternative Drosha cleavage is a general mechanism for isomiR production

Finally, we investigated the extent to which tertiary structure impacts the choice of Drosha cleavage sites for pri-miRNAs in general. To this end, we measured the Drosha cleavage fidelity of the top 200 miRNAs which are highly expressed in various tissues based on published miRNA-Seq results. We focused on 5’-arm-miRNAs (5p) since Drosha cleavages of these miRNAs may produce isomiRs with shifted seed sequences. The weighted average number of isomiRs with distinct 5’ ends was calculated for each individual miRNA (see *Materials and Methods* for detail) and used as a measurement of Drosha alternative cleavage. While the majority of the pri-miRNAs are processed precisely by Drosha, many pri-miRNAs have an alternative Drosha cleavage frequency higher than that of pri-miR-9-1 (Figure 5A). The cleavage profiles of each pri-miRNA are largely consistent among a wide range of tissues (18 in total), which supports the idea that an intrinsic feature such as tertiary structure is a major determinant for Drosha cleavage fidelity. However, a subset of pri-miRNAs displayed distinct Drosha cleavage profiles between tissues (Figure 5A), suggesting tissue-specific factors may also affect cleavage fidelity, possibly by modulating pri-miRNA structure.

All pri-miRNAs were ranked by the average number of 5’ isomiRs each produces (Figure 5A). The top 50 are classified as the low Drosha cleavage fidelity group whereas the bottom 50 are classified as the high-fidelity group. To analyze the lower stem length, we aligned all pri-miRNA sequences (miRBase v21) at the canonical Drosha cleavage site and calculated the percentage of paired bases at each position. The pri-miRNA basal junction can be inferred by a clear transition from unpaired (flanking sequence) to paired regions (lower stem). Consistent with previous studies (T. A. Nguyen et al., 2015), we observed that the average length of the lower stem, defined by the distance between the basal junction and the Drosha cleavage site, is 13nt on the 5p strand and 11nt on the 3p strand (Figure 5B). This result supports the current model in which the Drosha cleavage site is mostly determined by its distance to the basal junction (T. A. Nguyen et al., 2015). However, the same analysis of the low-fidelity group and the high-fidelity group revealed that both groups have better-defined basal junctions at expected positions compared with pri-miRNAs on average, suggesting the distance by itself is insufficient to determine Drosha cleavage fidelity.

In parallel, we tried to analyze 3D structures of pri-miRNAs in both groups. Since high-throughput approach in determining RNA tertiary structure is unavailable, we sought to first measure asymmetrical bulges on pri-miRNA stems, a 2D feature associated well with the stem distortion and flexibility in the case study of pri-miR-9 family. To this end, we calculated for each pri-miRNA an asymmetrical score, which is the sum of difference between numbers of nucleotides on each side of every bulge along the stem. Interestingly, while the average number of mismatches on pri-miRNA stem is similar (Figure 5–figure supplement 1A), the low fidelity group has a higher average asymmetrical score (Figure 5C), indicating that their stem structures are potentially more distorted and flexible. In line with this, most pri-miRNAs with low Drosha cleavage fidelity are predicted to have a distorted structure whereas the majority of the pri-miRNAs with high Drosha cleavage fidelity are relatively straight (Figure 5–figure supplement 1B). To measure their tertiary curvature quantitatively, we superimposed each structure with an ideal A-form RNA helix and calculated the root-mean-square deviation (RMSD) between their backbones (Figure 5D). A more distorted stem structure results in a larger RMSD value. As expected, pri-miRNAs in the low-fidelity group have a higher average RMSD than these of the high-fidelity group, implying that the overall distortion of a pri-miRNA stem impacts Drosha cleavage fidelity.

Together, these results demonstrate that the pri-miRNA tertiary structure, specifically the distorted stem correlates with alternative Drosha cleavage and production of isomiRs with altered seed sequences.

**Figure 5.**
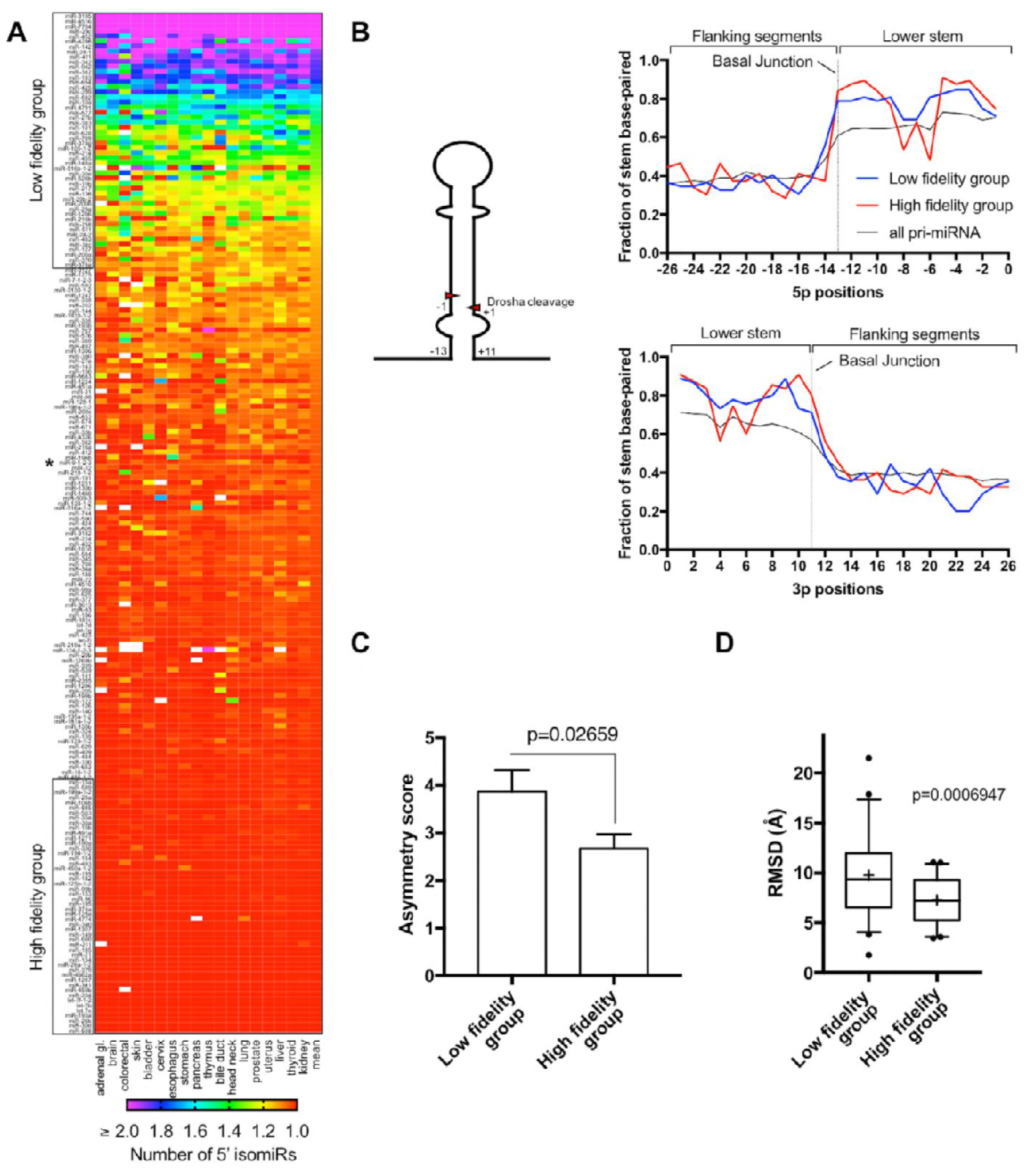
Pri-miRNA tertiary-structure-based alternative Drosha cleavage is a general mechanism for isomiR production. (***A***) Heatmap of the weighted average number of 5’ isomiRs generated from the top 200 5p miRNAs expressed in various tissues (TCGA normal tissue samples). White box indicates data unavailable. (***B***) Plot shows the fraction of pri-miRNAs that present a stem base-pairing on each position relative the Drosha cleavage site. Positions are numbered based on their distances to the Drosha cleavage site. Negative - upstream of the 5p cleavage site; positive - downstream of the 3p cleavage site. Black solid line shows the stem base-pairing of all the miRNA reported on miRBase21. (***C***) Plot of the average asymmetry score of pri-miRNAs with high (top 50) or low (bottom 50) Drosha cleavage fidelity. (***D***) Box-plot of the RMSD of pri-miRNAs with high (top 50) or low (bottom 50) Drosha cleavage fidelity.

## Discussion

The roles of RNase III enzymes in processing structured RNAs and in regulating gene expression are conserved from prokaryotic to eukaryotic cells (Court et al., 2013). Drosha, in particular, is essential for the maturation of most miRNAs. Understanding how Drosha precisely determines its cleavage sites is of great importance because this cleavage is the initial step in miRNA maturation. Downstream events, such as Dicer cleavage, RISC loading, post-maturation tailing/trimming and eventually miRNA target selection, all depend on the position where Drosha cleaves. Here, by combining deep sequencing and structural analysis, biochemical assays and functional studies, we demonstrated that pri-miRNA structure impacts Drosha cleavage fidelity. Importantly, we demonstrated that miR-9-alt, an isomiR resulting from an apparently low-frequency alternative Drosha cleavage, recognizes a distinct set of targets and is abundant in LGG, where it may function as a tumor suppressor. Thus, alterations in Drosha fidelity can profoundly influence cell function.

Previous models for Drosha cleavage focused on one-dimensional information (sequence motifs) and two-dimensional structural characteristics of pri-miRNAs. Here, for the first time, our results demonstrate that tertiary structure impacts Drosha cleavage fidelity. Given that RNase III enzymes in bacteria and yeast only accommodate substrates in the shape of A-form helices (Gan et al., 2006), it is reasonable to assume that Drosha has a similar preference based on its known structure (Kwon et al., 2016). Hence, pri-miRNAs with a bent or distorted stem in solution may need to alter its 3D conformation to fit in the Microprocessor complex. One possibility is that tension arising from this process induces Drosha cleavage at an alternative site. This model aligns well with previous studies of the other mammalian RNase III enzyme - Dicer: asymmetrical structural motifs in precursor hairpins, which are likely source of tertiary structure bending, induce Dicer alternative cleavages (Starega-Roslan et al., 2011). Alternatively, pri-miRNAs with distorted stems have higher flexibility, allowing them to fold into several distinct structures when complexing with Drosha. In this case, the alternative cleavage site may be a result of different configurations of the catalytic center and substrate. Future high-resolution structures of the ternary complex formed by Drosha, DGCR8 and pri-miRNA should give additional insights into the underlying mechanisms of Drosha alternative cleavage.

Although the existence of miR-9-alt was previously reported (Tan et al., 2014), we have both identified a novel role for this miRNA in LGG tumorigenesis and determined the pri-miRNA structural features that govern its biogenesis. In addition, we demonstrated that tertiary structure-based Drosha alternative cleavage is likely to be responsible for the generation of most, if not all, 5p isomiRs. Many factors are known to fine-tune Drosha cleavage efficiency (Ha and Kim, 2014; Jiang et al., 2017). Interestingly, we have shown that Drosha cleavage fidelity can vary between cell lines (Figure 1–figure supplement 1C) and tissues (Figure 5A), suggesting it is also subject to regulation, perhaps by cellular RNA-binding proteins. HnRNP A1 was reported to promote Drosha cleavage efficacy of pri-miR-18a by altering its structure (Michlewski et al., 2008). It is intriguing to hypothesize that other RNA binding proteins and helicases impact Drosha cleavage fidelity via modulating the folding of pri-miRNA. Future studies are required to understand how the cellular environment regulates Drosha cleavage fidelity and how the resulting isomiR changes affect cell physiology and disease development.

Although pri-miRNA paralogs are usually expected to function similarly, our demonstration that pri-mir-9-1 is primarily responsible for the production of miR-9-alt has revealed a new way in which individual paralogs can have distinct functions. It was proposed that during evolution, selective pressures stabilize alternative Drosha cleavage events leading to them becoming the dominant cutting site and resulting in the formation of new miRNAs (Berezikov, 2011). In support of this idea, Drosophila pri-miR-4 has high sequence homology to the Drosophila pri-miR-9 paralogs (9a, 9b and 9c) but has a different Drosha cleavage profile. The main cut site of the Drosophila pri-miR-4 is the same as the alternative Drosha cleavage site of human pri-miR-9-1, suggesting pri-miR-4 evolved from pri-miR-9 paralogs in flies. Thus, our work provides mechanistic insights into how miRNA paralog genes can be used as substrates to generate novel miRNAs. In this model of neofunctionalization, Drosha plays a central role in miRNA diversification and specificity.

Finally, the studies presented here have implications for RNAi technology, particularly for the design of shRNAs. A major source of off-target effects originates from heterogeneous products of shRNA processing *in vivo (Gu et al., 2012)*. We previously established the “loop-counting rule” of Dicer processing, which laid the groundwork for designing Pol III driven pre-miRNA-like shRNAs free of heterogeneous processing (Gu et al., 2012). Second generation pri-miRNA-like shRNAs, which can be expressed from a Pol II promoter and are thus more amenable to transcriptional control, require additional Drosha processing (Bofill-De Ros and Gu, 2016). Here, we demonstrated that imprecise Drosha processing drives the production of miRNA/siRNA with shifted seed sequences, generating undesired off-target repression. Hence, the tertiary structure of shRNA should be taken into consideration to avoid causing Drosha alternative cleavages. Our results provide additional guidelines for designing shRNAs with reduced off-target effects, which can then be used as tools for biological discovery and therapeutics.

## Materials and Methods

### Cell culture

HEK293T, HeLa cells and derived knockout cells were maintained in DMEM high glucose (Gibco) supplemented with 10% heat-inactivated fetal bovine serum (Hyclone), 100 U/ml penicillin-streptomycin (Gibco) at 37°C. Cells were tested to be free of mycoplasma contamination. Transfections were performed using PolyJet™ DNA Transfection Reagent (SignaGen) according to the manufacturer’s instructions.

### In vivo Drosha cleavage assay

Luciferase-based reporters were generated on the psiCHECK-2 vector (Promega). Pri-miR-9-1/-2/-3 inserts containing the pre-miRNA and ~200 nt flanking sequences at both ends were amplified by PCR from genomic DNA and inserted into the 3’UTR of the *Renilla* luciferase gene. Primers used in cloning are listed in Supplemental Table S2. 50 ng of the pri-miRNA reporter plasmids were co-transfected with either 50 ng of Drosha-expressing plasmids or empty vector in Drosha-KO cell lines. Cell lysates were obtained 48h post-transfection. Firefly and *Renilla* enzymatic activity were measured with Dual-Luciferase^®^ Reporter Assay System (Promega) and detected by GloMax^®^-Multi Luminescence Module (Promega) according to the manufacturer’s protocol. Microprocessor cleavage efficiency was calculated as follows:

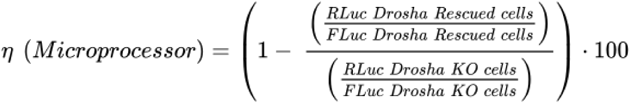
measured as the percentage of reporter activity in Drosha rescued to that where Drosha is knocked-out. Microprocessor efficiency was calculated as the complement of the Drosha cleavage ratio.

### In vitro Drosha cleavage assay

*In vitro* Drosha cleavage assay was performed similarly as previously described (T. A. Nguyen et al., 2015). In brief, FLAG-tagged Drosha and DGCR8 were co-expressed in HEK293T cells. At 48h post-transfection, Microprocessor complex was isolated by anti-FLAG immunoprécipitation. Pri-miRNA transcripts consisted of pre-miRNA and 200 nt flanking sequences on both sides were *in vitro* transcribed, purified, heat to 85°C and then refolded by slow cooling to 25°C for 15 minutes (ramp rate 0.1°C/sec). The microprocessor complex and pri-miRNAs were incubated at 37°C in reaction solution (6.4mM MgCl_2_ and 40U of RNase inhibitor) for 3h. Cleavage products were recovered using an acid phenol/chloroform extraction. Pre-miRNA products generated in the cleavage assay were detected by Northern blot.

### MicroRNA repression assay

Briefly, psiCHECK-2 reporters (Promega) with target sites for miR-9-alt or miR-9-can were inserted into the 3’UTR of the *Renilla* luciferase gene. Both strands of the target sequence were chemically synthesized, phosphorylated and ligated into the linearized psiCHECK-2 vector. Similarly, sh-miR-9-can/alt and sh-control were cloned downstream of a U6 promoter. Primers and oligonucleotides used in cloning are listed in Supplemental Table S2. 50 ng of the target reporter plasmids were co-transfected with either 50 ng of sh-miR-9-can/alt or sh-control into HEK293T cells. Cell lysates were obtained 48h post-transfection and measured with the Dual-Luciferase^®^ Reporter Assay System (Promega).

### Northern blot

Total RNA was isolated using Trizol (Life Technologies) and fractionated in 20% (w/v) acrylamide/8M urea gels. After, RNA was transferred to Hybond-N1 membranes (Amersham Pharmacia Biotech), crosslinked and blocked (PerfectHyb™ Plus Hybridization Buffer - Sigma). MicroRNA-9 (5p strand) was detected using ^32^P-labeled oligonucleotides. Images were obtained and analyzed using Amersham Typhoon (GE Healthcare).

### Small RNA NGS library preparation

5 μg of total RNA was ligated with RNA 3’ adaptor using T4 RNA Ligase 2 - truncated (NEB), in the presence of RNase Inhibitor (NEB). RNA 5’ adaptor was ligated using T4 RNA Ligase 1 - high concentration (NEB) and 10 mM ATP. Ligated small RNAs were reverse transcribed using superscript^®^ IV Reverse Transcriptase (Thermo-Fisher). Small RNA library cDNA was amplified and indexed using Phusion^®^ High-Fidelity DNA polymerase (NEB). Constructs were purified in a 6% (w/v) native acrylamide gel based on the expected product size and purified by ethanol precipitation. Library quality was assessed by using Qubit dsDNA HS Assay Kit (ThermoFisher) and Agilent High Sensitivity DNA kit (Agilent). Libraries were mixed together and prepared at a final concentration of 12pM and run on MiSeq Reagent Kit v3 (Illumina) according to the manufacturer’s specifications. Adaptors, primer sequences and detailed protocol temperatures can be found in Supplemental Table S2.

### Secondary and tertiary structure prediction

The sequences of pri-miR-9-1/-2/-3 was obtained from miRBase21 (Kozomara and Griffiths-Jones, 2014). Pri-miRNA sequences, excluding nucleotides upstream or downstream from the lower stem (Figure 4–figure supplement 1A) were subjected to folding with RNAstructure, using the default parameters for RNA to find the minimum free energy structure and close suboptimal solutions (Mathews et al., 2010). Predicted secondary structures were used as input to RNAComposer, a 3D structure prediction software (Popenda et al., 2012). 3D models were subjected to further refinements via molecular dynamics simulations with AMBER 14 package (Assisted Model Building with Energy Refinement). The AMBER force field ff14SB with ff99bsc0 and chi.OL3 parameter refinements for RNA were employed (Zgarbová et al., 2011). Implicit solvent simulations utilizing the Generalized Born model (GB) were performed, utilizing the latest corrections to the intrinsic Born radii parameters (mbondi3) in a GB-neck2 protocol (AMBER flag igb=8)(H. Nguyen et al., 2015; Tsui and Case, 2000). Simulations were run at 310 K, with a 2 fs time step and a Debye-Hückel (monovalent) salt screening concentration of 1.0. No cutoff was imposed on nonbonded interactions (cut = 999). The SHAKE algorithm was used to constrain all hydrogen bonds. The Langevin thermostat was employed with a collision frequency of 1.0 ps^−1^. A six-step 2.0 ns-long equilibration protocol was used that included energy minimization, heating to the target temperature of 310 K with harmonic restraints of 15 kcal/mol/Å^2^ on the RNA, followed by short MD stages with harmonic restraints gradually lowered from 10.0, down to 0.01 kcal/mol/Å^2^. Unrestrained (production) MD simulations were calculated every 2 femtoseconds for 250 ns. Post-processing of the MD trajectories (RMSD and average structure calculations) were generated with the cpptraj program of AMBER. Curvature of pri-miRNA tertiary structures was measured by calculating RMSD between pri-miRNA and a perfect A-form RNA helix using Pymol v2.0 (Schrodinger, LLC).

### SAXS sample preparation and data collection

Synthetic RNA (IDT) was used to synthesize exclusively the lower-stem, upper-stem and loop of pri-miR-9-1 (87nt) and pri-miR-9-2 (85nt). Concentration series SAXS measurements were carried out in order to remove the scattering contribution due to interparticle interactions and to extrapolate the data to infinite dilution. These RNAs were suspended in DEPC-treated water (Invitrogen) prepared at three concentrations (0.75, 1.2 and 1.8 mg/mL) in 50 mM HEPES buffer (GIBCO). The optimization of sample condition and screening of samples were performed on in-house SAXS instrument of NCI SAXS core. The selected samples were then measured at 12-ID-B beamline at the Advanced Photon Source in the Argonne National Laboratory. The procedures for data collection, processing, and analysis are similar to that previously described (Fang et al., 2013). The buffer background subtraction and intensity extrapolation to infinite dilution were carried out using NCI in-house developed MatLab script NCI-SAXS Core Facility.

### SAXS ensemble calculation

The ensemble calculation was performed using an NCI-SAXS-WAXS module deployed in the Xplor-NIH environment as published protocol (Schwieters and Clore, 2007). The NCI-SAXS-WAXS module allowed a simultaneous calculation of fitness between the experimental and back-calculated data in both SAXS regions. The equally sparse SAXS data, with q ranging from 0.004 to 0.89 Å^−1^ (a total of 72 data points), was used during the SAXS-restrained ensemble calculation. The difference in SAXS curves between experimental and back-calculated data is expressed as χ^2^, as defined:

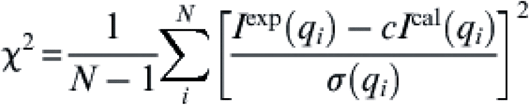
where c is a scaling factor, sigma(q_i_) is the experimental error, and I^exp^(q_i_) and I^cal^(q_i_) are the experimental and back-calculated scattering intensities of the i^th^ data point of the total N data points. A weighted harmonic energy potential function was used (E_SAXS_=C_SAXS_X^2^), where C_SAXS_ is a scaling factor. During the calculation, no Rg restraint was applied so that the ensemble outputs were allowed to freely sample the conformational space. Restraints were applied to maintain covalent geometry, prevent atomic overlap, and maintain the provided Watson-Crick and G:U wobble base-pairing in the duplexes. Knowledge-based restraints were applied to nucleic acid torsion angle conformations and to base-base packing (Kuszewski et al., 1997). The ensemble calculation was analyzed using an NCI-SAXS-WAXS data analysis module. All computation modules, scripts that contain all calculation parameters and conditions, and ensemble of the structures and SAXS data used for the calculation are provided upon request to the authors.

### MicroRNA expression and 5’ isomiR analysis

MicroRNA expression and 5’isomiR analysis were performed using QuagmiR on Amazon cloud instances through the Seven Bridges Genomics implementation of the NCI Cancer Genomics Cloud. QuagmiR is a customized Python scripts for the motif-based alignment and analysis of miRNA (https://github.com/Gu-Lab-RBL-NCI/quagmir). The analysis of The Cancer Genome Atlas (TCGA) was also performed using QuagmiR, with a previous conversion of the bam files to fastq files by Picard Sam-to-Fastq.

### Calculation of number of 5’ isomiRs

The weighted average number of 5’ isomiRs was calculated using an inverse Simpson index. This index measures the evenness of the 5’ isomiRs generated by each individual or family of paralog pri-miRNAs. The number of 5’ isomiRs (1/λ) was calculated as defined:

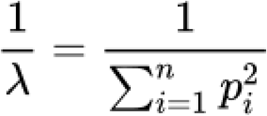
where n is each of the 5’ isomiRs detected and p is the weighted frequency of reads for that given 5’ isomiR in the sample. Normal tissue samples from TCGA (n=568) were used to generate the weighted average number of 5’ isomiRs in the corresponding tissue. Based on their average expression levels, we selected the top 200 most abundant miRNAs for heat-map plotting.

### Lower stem secondary structure analysis and asymmetry score

Genomic sequences of pri-miRNAs were obtained from UCSC Genome Browser. Each pri-miRNA analyzed consists of the pre-miRNA and flanking sequences of 30 nt on each side. The secondary structure was obtained (Gruber et al., 2008), and the bracket-dot notation of the lower stem was analyzed using custom R scripts (https://github.com/Gu-Lab-RBL-NCI). In brief, we first tested whether a nucleotide in one position is paired with another on the other side of the pre-miRNA. Nucleotides that failed such a test were labeled as unpaired. Then, we aligned all pri-miRNA 5p sequences by the 5’ Drosha cleavage site (5’ end of pre-miRNA) and aligned all 3p sequences by the 3’ Drosha cleavage site (3’ end of pre-miRNA). The fraction of paired nucleotides was calculated for each position and was plotted against its relative distance to the Drosha cleavage site. Similarly, we calculated an asymmetry score for each pri-miRNA. First, every bulge on the stem was analyzed individually and the absolute differences between numbers of nucleotides on each side was calculated. Asymmetry score was defined as the sum of this difference of all bulges along the pri-miRNA stem.

### Statistics

Statistical analysis was performed in GraphPad Prism7 statistical software. p values were calculated using t test, Mann-Whitney U-test or Wilcoxon test, as indicated. p value < 0.05 was considered statistically significant.

### Data Access

The Small RNA-Seq datasets generated in this study are available on NCBI GEO under the accession number GSE108893 (azefkigwdjephml - reviewer token). Previously published data sets used in this study are summarized in Supplemental Table S2. Other data are available upon reasonable request.

---

**Supplemental Table S1: Fold-change miR-9-alt predicted targets in LGG**

**Supplemental Table S2: Oligonucleotides and reference datasets used in this paper**

## Author contributions

X.B-DR. and S.G. designed all experiments and wrote the paper, with contributions from all co-authors. X.B-DR. conducted the experiments and data analysis with help from Q.C., M.J., L.D., A.Y. and T.-J.S. Structural analysis and MD simulations were done by W.K.K. and B.A.S. SAXS determination and ensemble analysis were done by Y.B., L.F. and Y-X.W.

## Supplementary Figure Legends

**Figure 1–figure supplement 1.** Pri-miR-9-1 has a unique Drosha cleavage profile. (***A***) Schematic representation of Drosha cleavage reporter assay. (***B***) CMV-driven pri-miR-9 cassettes were expressed individually in HeLa cells. pre-miR-9 and mature miR-9 were detected by Northern blot using probe-miR-9 (Supplemental Table S2). U6 snRNA was detected as an internal control. (***C***) Small RNAs from HeLa cells transfected with pri-miR-9-expressing plasmids were subjected to deep sequencing. Percentages of miR-9-can and miR-9-alt relative to all miR-9 reads were plotted. (***D***) Re-analysis of Drosha fCLIP (Kim, et al. Mol Cell 2017) from HEK293T cells where pri-miR-9-1 and pri-miR-9-2 is endogenously expressed. Percentages of canonical and alternative Drosha 5’ cleavage are depicted together with preferred Drosha 3’ cleavage site.

**Figure 2–figure supplement 1.** miR-9-alt regulates a novel set of targets to repress low grade glioma (LGG). (***A***) Left, schematic representation of the shRNAs: sh-miR-9-can and sh-miR-9-alt. Encoded miR-9-can or miR-9-alt is in red and the corresponding seed is boxed. The scheme also illustrates the disruption of the seed of the miR-9-3p strand. Right, sh-miR-9-can and sh-miR-9-alt were expressed individually in HEK293T cells. Small RNAs were measured by deep sequencing. Right, quantification of miR-9-can and miR-9-alt is listed. (***B***) Target sites of either miR-9-can or miR-9-alt were cloned individually in the 3’UTR of the luciferase reporter. Dual-luciferase assays were performed at 48h post-transfection, and the results were plotted as described previously. Error bars represent the SEM from three biological replicates. (***C***) Heatmap of the most abundant miRNAs in each tumor type reported on TCGA. (***D***) Diagram of the analysis pipeline used to obtain the Cumulative curves in Figure 4D. (***E***) Levels of pri-miR-9 paralogs transcripts were measured using Fragments Per Kilobase of transcript per Million mapped reads (FPKM) of the corresponding host gene in LGG patient samples (n=525). (***F***) Kaplan-Meier survival curve of LGG patients with high (red line) and low (black line) levels of the pri-miR-9-2 host gene (p=0.17, non-significant). p-values were calculated using Wilcoxon test. (***G***) GRIN2A (Glutamate Ionotropic Receptor NMDA Type Subunit 2A-NMDA2A) a predicted target for miR-9-alt. Predicted target site was cloned in the 3’UTR of the luciferase reporter. Dual-luciferase assays were performed at 48h post-transfection, and the results were plotted as described previously. Error bars represent the SEM from three biological replicates. **, p < 0.01 n.s., non-significant.

**Figure 3–figure supplement 1.** The lower stem plays a major role in determining cleavage fidelity. (***A***) Microprocessor efficiency on pri-miR-9 chimeras was measured by a Drosha cleavage reporter assay. Results were plotted as described in Figure 1A. (***B***) CMV-driven pri-miR-9 chimeras were expressed individually in HEK293T and HeLa cells. pre-miR-9 and mature miR-9 were detected by Northern blotting with probe-miR-9. (***C,D***) Sequence and secondary structure of the pri-miR-9 chimeric constructs. Green labeling illustrates the nucleotides changed in each case. The frequency of cleavage at each position in HEK293T cells is indicated. (***E,F***) Relative frequency of Drosha alternative cleavage in each pri-miR-9 chimeric construct compared to that of pri-miR-9-1 in HeLa cells. (***G***) Sequence alignment of pri-miR-9 paralogs in vertebrates performed using ClustalO. Conserved nucleotides are labeled with asterisk and variable nucleotides are labeled in red.

**Figure 4–figure supplement 1.** The tertiary structure of the lower stem drives alternative Drosha cleavage. (***A***) Secondary structure of pri-miR-9-1, −2 and −3, predicated by RNAStructure. (***B***) Three-dimensional models (5p and 3p strands are colored red and lower stem in cyan). RMSD fit of 3D molecular dynamics structures to an ideal A-form RNA helix over the course of the simulation. RMSD was calculated from the pri-miR-9 paralog structures (excluding nucleotides beyond the basal junction) before and after the molecular dynamics simulation. (***C***) From left to right. Chi-square values obtained from 100 structures generated in the ensemble calculation (Ne) from size 1 to 8. Detailed view of the Intensity(q) plot of pri-miR-9-1 and −2 at at q>0.2 Å^−1^ and Ne=1. Distribution of residuals along the q for the top 10 ensemble calculations. (***D***) Northern blot validating the expression of mature miR-9 from pri-miR-9-1-a and -b in HEK293T and HeLa cells. (***E***) Table containing the frequency of Drosha cleavage at each position in HEK293T and HeLa cells. (***F***) Secondary structure of pri-miR-9-1 lower-stem-corrected mutants, pri-miR-9-1-perfect-stem and pri-miR-9-1-bulged-stem, and expected basal junction. (***G***) RMSD fit of 3D molecular dynamics structures to an ideal A-form RNA helix over the course of the simulation of pri-miR-9-1-perfect-stem and pri-miR-9-1-bulged-stem. (***H***) Northern blot validating the expression of mature miR-9 from pri-miR-9-1 lower-stem-corrected mutants in HEK293T and HeLa cells. (***I***) Frequency of Drosha alternative cleavage on each of the pri-miR-9-1 corrected mutant constructs relative to that of wild-type pri-miR-9-1 was measured in HEK293T and HeLa cells by deep sequencing and plotted here.

**Figure 5–figure supplement 1.** Pri-miRNA tertiary-structure-based alternative Drosha cleavage is a general mechanism for isomiR production. (***A***) Plot of the average number (#) of unpaired nucleotides in the stem of pri-miRNAs with high (top 50) or low (bottom 50) Drosha cleavage fidelity. (***B***) Examples of the predicted tertiary structures of pri-miRNAs in the low-fidelity group.

